# YAP1 induces invadopodia formation by transcriptionally activating TIAM1 through its enhancer in breast cancer

**DOI:** 10.1101/2021.03.02.431738

**Authors:** Jie Shen, Qingwen Huang, Shengjie Feng, Weiyi Jia, Liang Liu, Xiaolan Li, Deding Tao, Daxing Xie

**Author notes:** These authors contributed equally to this work (Jie Shen, Qingwen Huang). Correspondence: Daxing Xie, M.D., Ph.D. 1095 Jiefang Ave. Wuhan, Hubei 430030, China. **Funding:** This work was supported by grants from the National Natural Science Foundation of China, Grant numbers: 81572861, 81773053. **Conflict of Interest:** The authors have no conflicts of interest to disclose.

## Abstract

Yes-associated protein 1 (YAP1), a central component of the Hippo pathway, plays an important role in tumor metastasis; however, the mechanism remains to be elucidated. Here, we reported that YAP1 could induce invadopodia formation and promote tumor metastasis in breast cancer cells. We identified that TIAM1, the guanine nucleotide exchange factor, is a target of YAP1-TEAD4 complex. YAP1 promotes TEAD4 binding to the enhancer region of TIAM1, which activates TIAM1 expression and subsequently increases RAC1 activity. These findings reveal the functional role of Hippo signaling in invadopodia, and provide potential molecular targets for preventing tumor metastasis in breast cancer.

**Significance:** Through regulating the enhancer region of TIAM1, YAP1 induces invadopodia formation and promotes tumor metastasis in breast cancer.

## Introduction

Breast cancer is the most common malignant disease in women and causes severe cancer-related death (1). Although great improvements have been achieved in tumor screening and systematic therapy, metastasis remains the main cause of cancer lethality and a major obstacle in the treatment of breast cancer (2, 3). Metastasis contains a series of sequential interrelated steps, including detachment from the primary lesion, invasion, extravasation and establishment of a microenvironment. Numerous cellular signaling pathways and molecules are involved in these processes (4).

Recently, Hippo signaling pathway has been reported to be strongly involved in cancer progression and plays an important role in tumor metastasis (5). Hippo signaling was first reported 20 years ago in *Drosophila* as a regulator of cell proliferation and apoptosis (6). Subsequent studies have revealed its high conservation and complex functions in mammals (6–8). In recent years, the Hippo pathway has emerged as a central player in many aspects of tumor biology (9–11). The transcriptional coactivator YAP1 (yes-associated protein 1), which is also known as YAP or YAP65, is the main component of the Hippo pathway. After Hippo signal is inhibited, YAP is dephosphorylated at serine 127 and transferred into the nucleus where it induces target genes expression via binding with related transcription factors, especially TEA domain family members (TEADs) (12–15). YAP is reported to function as an oncogene, and its hyperactivation leads to a variety of tumor-promoting effects (16, 17). Previous studies indicated that YAP overexpression is related with breast and brain tumor metastasis (18). In addition, YAP overexpression can trigger the epithelial-mesenchymal transition (EMT) (17, 19, 20) and regulate actin dynamics (21, 22). These findings implied the critical role of YAP in promoting tumor cell metastasis; however, the mechanism remains to be elucidated.

Invadopodia are actin-rich protrusions of the basolateral plasma membrane and are associated with degradation of the extracellular matrix in cancer invasiveness and metastasis (23–26). Previous studies have revealed that invadopodia are composed of structural proteins, such as cortactin, TKS4 and TKS5 (27–29), and are regulated by numerous regulatory proteins that control actin dynamics (30). EMT-related genes, such as TWIST1, and Rho GTPases, such as RAC/CDC42, have also been reported to play important roles in invadopodia formation (26, 31–34).

In this study, we reported that YAP1, interacting with transcriptional factor TEAD4, induces invadopodia formation and promotes tumor metastasis in breast cancer cells. We identified that the TIAM1, guanine nucleotide exchange factor, functions as a target of YAP1-TEAD4 complex. YAP1 promotes TEAD4 binding to the enhancer region of TIAM1, which up- regulates TIAM1 expression and subsequently increases RAC1 kinase activity. These findings revealed the novel role of Hippo signaling pathway in invadopodia, and provided potential molecular targets for preventing tumor metastasis in breast cancer.

## Results

### YAP1 induces invadopodia formation and is associated with tumor metastasis in breast cancer

To evaluate the role of YAP1 in invasive breast cancer (IBC), we collected clinical specimens from 6 normal/benign breast disease tissues, 21 primary tumors, 6 positive lymph nodes and 9 distant metastatic lesions, and detected YAP1 expression by IHC staining. As shown in Figure 1A, YAP1 protein was over-expressed in IBC. Compared with its expression in primary tumor, YAP1 protein levels were significantly increased in positive lymph nodes and distant metastatic lesions, whereas YAP1 protein was mainly localized in the nucleus (Fig. 1A, general characteristics of patients are presented in Datasets S1). To determine a potential relationship between YAP1 signaling and tumor metastasis in IBC, we performed gene set enrichment analysis (GSEA) of the expression profile of purified tumor cells from 14 primary breast tumors and 6 metastatic lymph nodes available from the GSE database (GSE30480). GSEA data showed that the YAP-conserved gene signature was positively associated with lymphatic metastasis (Fig. 1B and Datasets S2). Prognosis data indicated that patients with YAP1-high expression had worse disease-free survival than those with YAP1-low expression based on two different datasets (Fig. 1C. GSE21653, n=252, p=0.021 and GSE25006, n=508, p<0.01). To further determine whether nuclear localization of YAP1 was associated with poor prognosis, we analyzed YAP1-regulated downstream genes expression, including AMOTL2, CYR61, CTGF, MYC, AREG, GLI2 and AXL (35, 36), in different prognosis groups of IBC patients from the TCGA dataset (n=962) based on the SurExpress program (37). Our results showed that the expression levels of YAP1, as well as its target genes (AMOTL2, CYR61, CTGF, MYC, GLI2 and AXL) increased significantly in the poor prognostic group of IBC patients (Fig. S1A, p<0.01). Taken together, these results demonstrated that overexpression and nuclear localization of YAP is associated with tumor invasiveness and poor prognosis in IBC patients. To evaluate the functional role of YAP1 on tumor metastasis, we examined its endogenous expression in 4 breast cancer cell lines (MCF7, T47D, MDA-MB-231 and MDA-MD-468) and one nontumorigenic breast epithelial cell line (MCF10A). Western blot results showed that YAP1 protein was relatively high-expressed in T47D, MDA-MB-231 and MDA-MD-468 cell lines, and low-expressed in MCF7 and MCF10A cell lines (Fig. 1D). Given that MCF-7 and MDA-MB-231 cells exhibited low and high metastatic potential, respectively (38), these two cell lines were chosen for further studies. In MCF7, overexpression of YAP1 wild-type (WT) protein was transfected with pcDNA3.1-YAP1 plasmid (Fig. 1E), in contrast, a collection of siRNAs was used to knockdown endogenous YAP1 expression in MDA-MB-231 cell line (Fig. S1B). Transwell migration, invasion assays, and wound-healing assay showed that YAP1 can regulate breast cancer cell migration and invasion in vitro (Figures S1C, D, E, F), which are consistent with our previous results (39).

**Figure 1.**
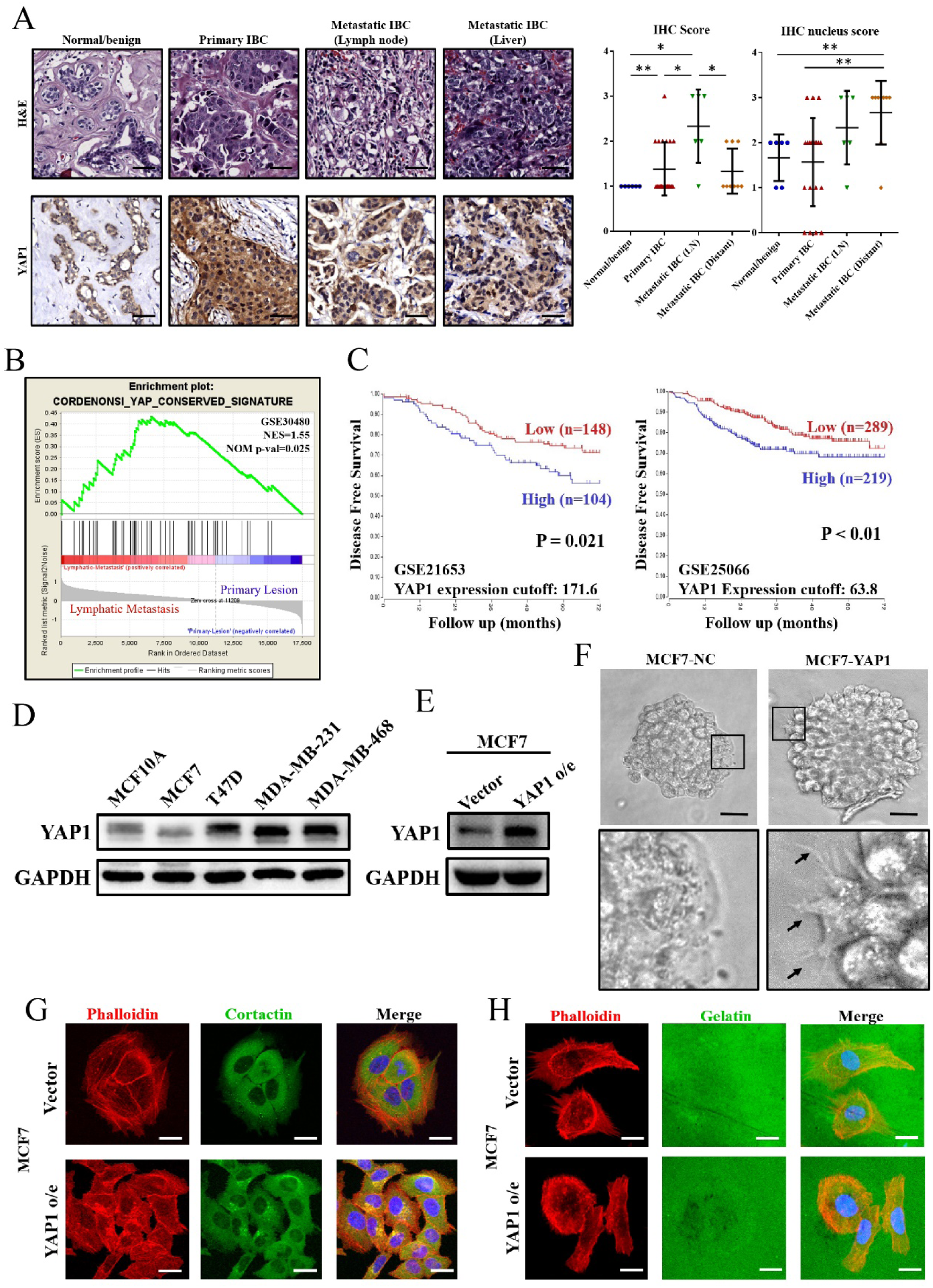
**YAP1 is associated with tumor invasiveness and induces invadopodia formation in breast cancer**. (A) YAP1 expression according to breast cancer sampling site. IHC score of YAP1 was significantly increased in IBC compared with normal/benign breast disease tissue. In IBC specimens stratified by sampling sites, YAP1 exhibited relatively high expression in metastatic lymph nodes, whereas YAP1 nuclear expression was significantly increased in distant metastatic lesions. (H&E: hematoxylin-eosin staining, IHC: immunohistochemistry, IBC: invasive breast cancer, LN: lymph node, *p<0.05 and **p<0.01 by Student’s t test, Scale bar: 50 um). (B) Gene set enrichment analysis (GSEA) of purified tumor cells from 14 primary breast tumor tissues and 6 metastatic lymph nodes from the GEO database (GSE30480). Representative GSEA plots indicated that the conserved YAP signature was positively associated with lymphatic metastasis among 189 oncogenic signature gene sets (NES=1.55, NOM p-value < 0.05). (C) Kaplan-Meier disease-free survival analysis based on YAP1 mRNA expression in 252 breast cancer patients (GSE21653) (Left) and 508 breast cancer patients (GSE25066) (Right). YAP1 overexpression was associated with a high risk of recurrence. Analysis was based on R2: Genomics Analysis and Visualization Platform. (D) YAP1 was frequently upregulated in highly invasive breast cancer cell lines as determined by Western blot. Endogenous YAP1 was relatively high expression in MDA-MB-231 cell line and was low expression in MCF7 cell line. (E) Western blot verified overexpression of YAP1 in MCF7 cell line. (F) YAP1 overexpression in MCF7 cells promoted pseudopod-like structure (black arrows) formation in 3D tumor spheroid invasion assays (Scale bar: 40 µm). (G) YAP1 overexpression promoted invadopodia formation in MCF7 cells. Invadopodia were visualized by colocalization of cortactin (green) and F-actin (stained with phalloidin, red). Scale bar: 20 µm. (H) YAP1 promoted ECM degradation ability of MCF7 cells. Cells were plated on Alexa Fluor 488-conjugated gelatin (green) for 24 hr. F-actin was stained with phalloidin (red), and nuclei were stained with DAPI (blue). The area of gelatin degradation appears as a black area beneath the cells (scale bar: 20 µm).

To further mimic the invasiveness of breast cancer cells in the microenvironment, we performed 3D tumor spheroid invasion assays (40). Interestingly, we observed that YAP1 overexpression induced the formation of pseudopod-like structures on the rim of tumor spheres in MCF7 cells (Fig. 1F). Invadopodia, known as the structure with cancer-specific pseudopodium, could be detected by co-localization of F-actin with the actin-bounding protein cortactin through immunofluorescence assay (41). Thus, we determined the effect of YAP1 on invadopodia formation in vitro. Our results showed that YAP1 overexpression induced invadopodia formation in MCF7 cells (Fig. 1G), whereas knockdown of endogenous YAP1 significantly inhibited invadopodia formation in MDA-MB-231 cells (Fig. S1G). To detect the ECM degradation ability of invadopodia, we performed gelatin degradation assay. Our results showed that YAP1 overexpression promoted gelatin degradation in MCF7 cells significantly (Fig. 1H), whereas knockdown of YAP1 decreased the gelatin degradation ability of MDA-MB-231 (Fig. S1H). Taken together, our results demonstrated that YAP1 could regulate invadopodia formation in breast cancer cells.

### YAP1-TEAD4 interaction is essential for invadopodia formation

YAP1 is a transcriptional coactivator that trans-locates into the nucleus after dephosphorylation of serine 127. After that, it combines with TEADs though the TEAD-binding domain or SMADs though the WW domain to activate downstream genes expression (42). To determine the functional domain of YAP1 involved in invadopodia regulation, we transfected YAP1 constitutive nuclear-localized mutant (Flag-YAP1-S127A) or TEAD-binding domain mutant (GFP-YAP1-S94A) into MCF7 cell lines (Fig. 2A, 2B). Compared to YAP1-S94A, the YAP1-S127A mutant exhibited increased expression level of TEAD target gene CTGF and CYR61 (Fig. S2A). Then, we determined whether the YAP1-S127A or YAP1-S94A mutant could induce invadopodia formation. Using immunofluorescence, we observed that ectopic expression of the YAP1-S127A mutant, rather than YAP1-S94A, induced invadopodia formation in MCF7 cells (Fig. 2C). To further determine the role of the YAP1-TEAD interaction in invadopodia formation, we used verteporfin, a small molecular inhibitor of the YAP-TEAD interaction (43). Immunofluorescence and gelatin degradation assays showed that, after being treated with verteporfin (10 µM), either MCF7-YAP1-S127A or MDA-MB-231 (expressing endogenous YAP1) cells exhibited decreased invadopodia formation (Fig. 2D, Fig S2B) and ECM degradation ability significantly (Fig. 2E, Fig. S2C). In addition, Transwell assays demonstrated that verteporfin inhibited cell migration in both MCF7-YAP1-S127A and MDA-MB-231 cells (Fig. S2D). 3D tumor spheroid invasion assay showed that verteporfin treatment decreased pseudopod-like structures in MCF7-YAP1 cells (Fig. S2E). Thus, our results suggested a critical role of the interaction between YAP and TEAD family proteins in invadopodia formation.

**Figure 2.**
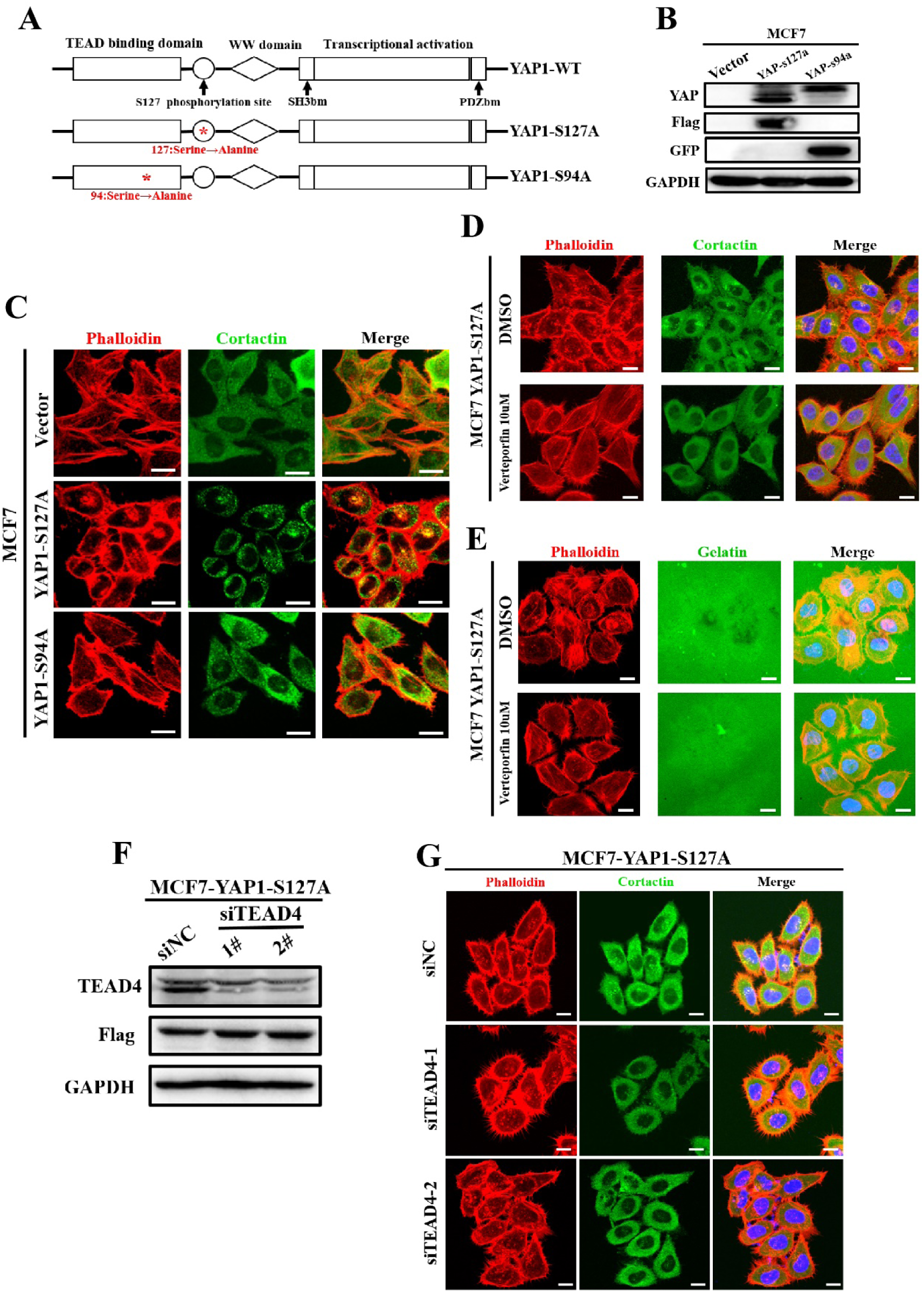
YAP1-TEAD4 interaction is essential for invadopodia formation and tumor migration. (A) Domain organization of YAP1 wild type (YAP1-WT), YAP1 constitutively activated mutant (YAP1-S127A) and YAP1 TEAD-binding domain mutant (YAP1- S94A). Red asterisk indicates the mutation site. (B)YAP1 protein expression in MCF7 cells following transfection with pcDNA3.1-YAP1-S127A (flag-tagged) and pcDNA3.1-YAP1-S94A (GFP-tagged). (C) Ectopic expression of YAP1-S127A not YAP1-S94A promoted invadopodia formation in MCF7 cells. (Actin: red, stained with phalloidin, Cortactin: green, stained with DyLight 649; scale bar: 20 µm). (D) Verteporfin inhibited invadopodia formation in MCF7 cells expressing YAP1- S127A mutant. Cells were seeded on 0.1% gelatin and treated with 10 µM verteporfin (DMSO was used as negative control). After 48 h of culture, invadopodia were visualized by colocalization of cortactin (green) and F-actin (red). Scale bar: 20 µm. (E) Verteporfin inhibited ECM degradation in MCF7 cells transfected with YAP1- S127A mutant. MCF7 overexpressing YAP1-S127A mutant plasmid were seeded on Alexa Fluor 488-conjugated gelatin (green) and treated with 10 µM verteporfin (DMSO was used as negative control) for 24 hr. F-actin was stained with phalloidin (red), and nuclei were stained with DAPI (blue). The area of gelatin degradation appears as black area beneath the cells (scale bar: 20 um). (F) After transfection with YAP1-S127A (flag-tagged) plasmid, siRNAs were used to knockdown endogenous TEAD4 expression in MCF7 cell line. (G) Knockdown of endogenous TEAD4 expression reversed YAP1-S127A mediated invadopodia formation in MCF7 cell line. Scale bar: 20 µm.

The TEAD family proteins include four highly homologous members (TEAD1, TEAD2, TEAD3, and TEAD4) that exhibit similar biological function (44, 45). To determine the functional TEAD protein for invadopodia formation, we examined expression levels of TEAD1-4 in MCF7 cells. Our results showed that TEAD1 and TEAD4 mRNA levels were high-expressed, whereas TEAD2 and TEAD3 mRNA could be hardly detected in MCF7 cells (Fig. S2F). Next, we used the cBioPortal program to assess TEAD expression in clinical breast cancer specimens. We found that TEAD4 exhibited the highest levels of up-regulation among the TEAD family members in breast cancer (Fig. S2G). In addition, SurvExpress analysis revealed that TEAD4, not TEAD1 expression exhibited a positive correlation with poor prognosis in breast cancer patients (Fig. S2H). Therefore, we suggested TEAD4 is the candidate for YAP1-TEAD complex in the study. Our further results showed that knocking down endogenous TEAD4 expression could significantly reverse YAP1-S127A induced invadopodia formation and cell migration (Fig. 2F, 2G and Fig. S2I). Similarly, combination of knocking down YAP1 and/or TEAD4 expression could further inhibited invadopodia formation and cell migration in MDA-MB-231 cells (Figures S2I, S2J, S2K). These results demonstrated the YAP1-TEAD4 interaction is essential for invadopodia formation in breast cancer cells.

### YAP1-TEAD4 regulates invadopodia formation in a TIAM1-RAC1 dependent manner

To identify the functional downstream of YAP1-TEAD4, we extracted mRNA from MCF7 cells overexpressing either YAP1-S127A or control plasmid, and profiled gene expression using the cDNA microarray (Datasets S3). Then, these different expressed genes (genes with at least a 1.5-fold change in expression after YAP1-S127A induction) were matched with identified TEAD4-binding gene from ENCODE ChIP-sequence dataset GSM1010860 (Fig. 3A). Based on this criterion, we screened out 272 up-regulated TEAD4-binding genes and 214 down-regulated TEAD4- binding genes in MCF7 cells (Fig. 3B upper). To explore the major function of these YAP1-TEAD4 binding genes, gene ontology (GO) enrichment analysis was performed (Fig. S3). Our analysis implied that “Rho/Rac guanyl-nucleotide exchange factor (GEF) activity” was included in the top 5 GO enrichment (molecular function) categories of up-regulated TEAD4- binding genes (Fig. 3B lower). These data suggested YAP1-TEAD4 complex might regulate Rho/Rac family activity.

**Figure 3.**
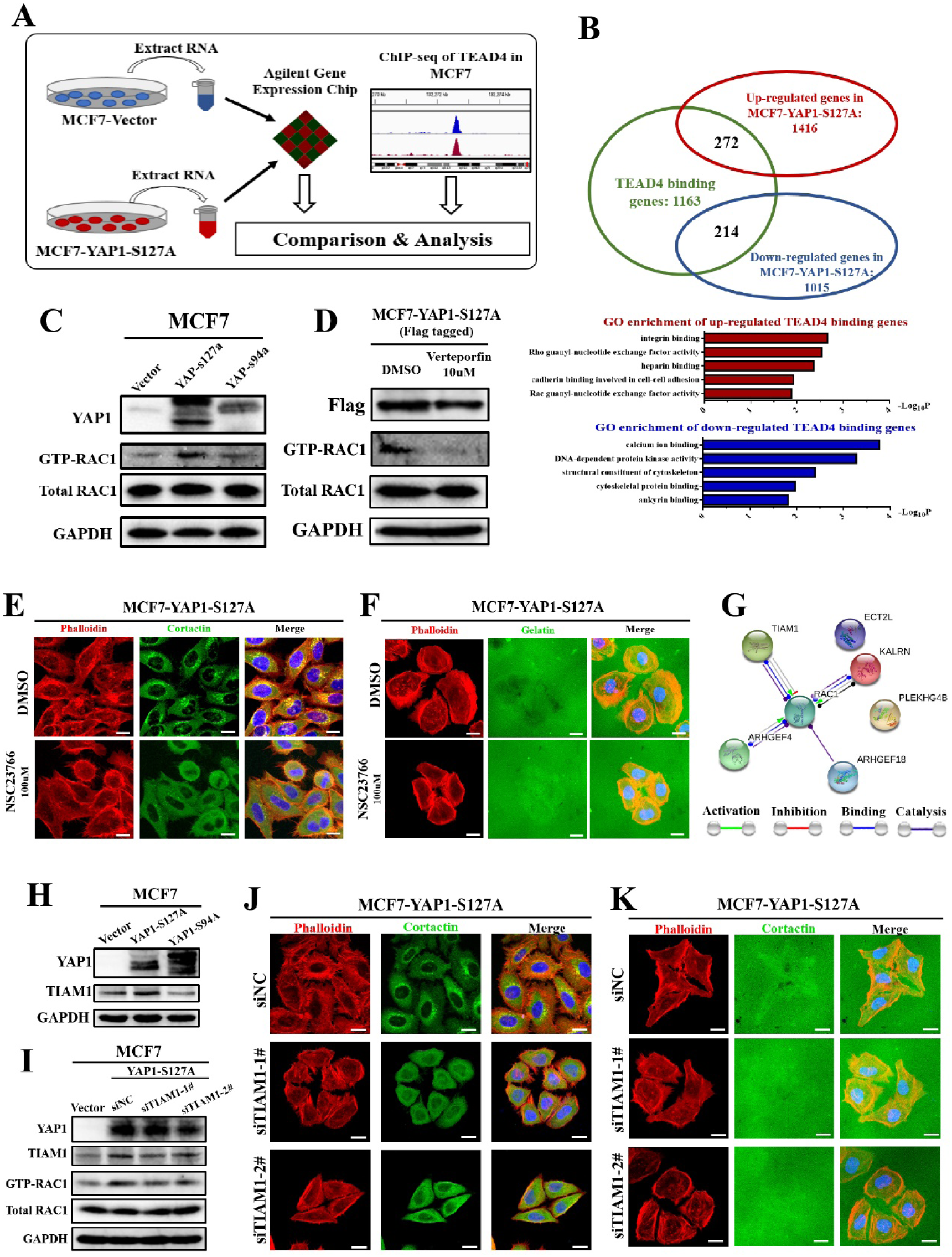
YAP1-TEAD4 regulates invadopodia formation in a TIAM1-RAC1 dependent manner. (A) Experimental design to determine transcriptional targets of the YAP1-TEAD4 complex in MCF7 cells. MCF7 cells were transfected with vector and YAP1-S127A mutant for 48h. Then, total RNA was subjected to expression profiling using the Agilent SurePrint G3 Human Gene Expression v3 Panel (8*60K, Design ID: 072363). The profiling results were matched with ChIP-seq data of TEAD4 in MCF7 cells from the ENCODE database (GSM1010860). (B) Candidate TEAD4-binding (TB) genes regulated by YAP1-S127A in MCF7 cells. TB genes acquired from ENCODE were analyzed, including 272 upregulated genes and 214 downregulated genes in MCF7-YAP1-S127A cells. These up- and down- regulated TEAD4-binding genes in were used in Gene Ontology enrichment analysis. Top 5 biological processes involved in molecular function are presented. Bar plot representation of Log10P values. (C) Expression of YAP1-S127A not YAP1-S94A promoted RAC1 activation in MCF7 cells as determined by GTP-bound GTPase pull-down experiments and Western blots. (D) Verteporfin reversed YAP1-S127A-induced RAC1 activation in MCF7 cells. MCF7 cells (transfected with the flag-tagged-YAP1-S127A overexpression plasmid) were treated with 10 µM Verteporfin (DMSO was used as negative control) for 24 hr. Then, RAC1 activation was measured by GTP-bound GTPase pull-down experiments and Western blots. (E) NSC23766 reversed invadopodia formation, which was induced by the YAP1- S127A mutant in MCF7 cells. MCF7 cells (expressing the YAP1-S127A overexpression plasmid) were seeded on 0.1% gelatin and treated with 100 µM NSC23766 (DMSO was used as a negative control). After 24 h of culture, invadopodia were visualized by colocalization of cortactin (green) and F-actin (red). Nucleus was strained with DAPI (blue). Scale bar: 20 µm. (F) NSC23766 reduced ECM degradation in MCF7 cells transfected with YAP1-S127A mutant. MCF7 cells (expressing YAP1-S127A overexpression plasmid) were seeded on Alexa Fluor 488-conjugated gelatin (green) and treated with 100 µM NSC23766 (DMSO was used as a negative control) for 24 hr. F-actin was stained with phalloidin (red), and nuclei were stained with DAPI (blue). The area of gelatin degradation appears as a black area beneath the cells (scale bar: 20 um). (G) STRING analysis was used to identify the interaction between RAC1 and the upregulated TEAD4-binding genes from Rho/RAC guanyl-nucleotide exchange factor activity categories (GO: 0005089 and GO: 0030676) in MCF7. TIAM1, ARHGEF4 and KALRN presented the potential activators of RAC1. (H) YAP1-S127A not YAP1-S94A increased TIAM1 protein levels in MCF7 cells as determined by Western blot. (I) After transfection with the YAP1-S127A mutant, TIAM1 knockdown reversed RAC1 activation in MCF7 cells. (J) Knockdown of TIAM1 expression reversed YAP1-induced invadopodia formation in MCF7 cells. Green: cortactin; red: F-actin; blue: DAPI. Scale bar: 20 µm. (K) TIAM1 knockdown reversed YAP1-induced ECM degradation in MCF7 cells. Green: gelatin; red: F-actin; blue: DAPI. Scale bar: 20 µm.

Previous studies reported that RAC1 played a critical role for invasive protrusion formation (46) and ECM degradation (47). RAC1 activity was required for cortactin tyrosine phosphorylation which is a crucial step for invadopodia assembly (48). To determine RAC1 kinase activity, we performed GST-pull-down assays to detect GTP-bound RAC1. Our data demonstrated that compared with the YAP1-S94A mutant, YAP1-S127A overexpression significantly increased RAC1 kinase activity in MCF7 cells. Meanwhile, knocking down YAP1 endogenous expression or using verteporfin could inhibit RAC1 activation in both MCF7-YAP1-S127A and MDA-MB-231 cells (Fig. 3C, 3D and Fig. S4A, S4B). CDC42 is also a small GTPase involved in invadopodia formation (34). Our data showed that CDC42 activity was not affected in MCF7 cells expressing YAP1- S127A mutant (Fig. S4C). To further determine the role of endogenous RAC1 in YAP1 regulated invadopodia function, we treated YAP1-S127A mutant expressing MCF7 cells with NSC23766, a RAC1 inhibitor, or RAC1-siRNA (Fig. S4D, S4E). Both NSC23766 and RAC1-siRNA treatment could block YAP1-induced invadopodia formation (Fig 3E, Fig. S4F) and cell migration (Fig. S4G, S4H). NSC23766 could also reverse YAP1-induced ECM degradation in MCF7 cell line (Fig. 3F). Similarly, inhibiting of RAC1 activity also reduced migration, invadopodia formation and ECM degradation in MDA-MB-231 cells (Fig. S4G, S4I, S4J). These data clearly demonstrated that RAC1 is critical for YAP1 regulated invadopodia formation and tumor migration.

To further elucidate the mechanism of YAP1-induced RAC1 activation, we analyzed the YAP1-S127A up-regulated TEAD4-binding genes, and used heat-map to summarize 6 candidate genes classified in the Rho/Rac GEF activation GO category (Fig. S5A). By using the String program (string- db.org), we identified TIAM1, ARHGEF4 and KALRN as potential activators of RAC1 (Fig. 3G). After further validating their mRNA expression through RT-qPCR, we concluded that TIAM1 was the most significant up-regulated gene in MCF7 cells overexpressing YAP1-S127A (Fig. S5B). Consistently, YAP1-S127A overexpressing significantly increased TIAM1 protein levels in MCF7 cells, whereas the YAP1-S94A mutant could not (Fig. 3H). In addition, verteporfin could reverse TIAM1 expression in MCF7 cells overexpressing YAP1-S127A (Fig. S5C). Similarly, either knocking down endogenous YAP1 or inhibition of the YAP1-TEAD interaction could reduce TIAM1 protein expression in MDA-MB-231 cells (Fig. S5D, S5E). To confirm the role of TIAM1 in YAP1- TEAD4 regulated RAC1 activation and invadopodia formation, we knocked down endogenous TIAM1 expression in MCF7 cells overexpressing YAP1-S127A mutant and detected GTP-bound RAC1 via GST-pull down assay. Our result showed that knocking down endogenous TIAM1 reversed YAP1-induced RAC1 activation (Fig. 3I). In addition, TIAM1 knockdown significantly reduced invadopodia formation (Fig. 3J), ECM degradation (Fig. 3K) and migration ability in MCF7 cells overexpressing YAP1-S127A mutant (Fig. S5F). Finally, we performed gene correlation analysis based on 107 breast cancer patients in Auckland (GSE36771) and 1097 breast cancer patients from TCGA. Our analysis revealed that TIAM1 gene expression levels were positively correlated to YAP1 levels (n=107, R=0.408, p<0.001 in Auckland and n=1097, R=0.177, p<0.001 in TCGA) in clinical breast cancer specimens (Fig. S5G). Taken together, our results indicated that TIAM1 plays an important role in YAP1-TEAD4 regulated RAC1 activation and invadopodia formation.

### YAP1-TEAD4 transcriptionally activates TIAM1 expression through its enhancer region

Through binding to the promoter/enhancer region of targets genes, the YAP-TEAD complexes increase genes transcriptional activity (49–51). To characterize the transcriptional regulation of TIAM1 via the YAP1-TEAD4 complex, we annotated TEAD4 ChIP sequence data of MCF7 cell line from ENCODE database via ChIP-Seek software (http://chipseek.cgu.edu.tw/). Interestingly, we found that the TEAD4-binding site (TBS) was not located in the traditional promoter region of TIAM1 (Datasets S4). Instead, it was located in the binding region of H3K27ac where is an active enhancer marker (Fig. 4A and Fig. S6A). Our ChIP results clearly demonstrated that overexpressing of YAP1-S127A mutant could promote YAP1-TEAD4 complex binding to the TBS region of TIAM genome in MCF7 cells (Fig. 4B and Fig. S6B). To further determine its transcriptional activity, we constructed a luciferase-based reporter containing the TBS region (pGL3- TBS-Luc). Dual luciferase reporter assay demonstrated that the YAP1- S127A could significantly promote TBS-regulated luciferase activity in both HEK293T and MCF7 cells (Fig. 4C), where YAP1-S94A mutant could not (Fig. S6C). These results suggested that YAP1-TEAD4 complex might active TIAM gene through binding to its enhancer region.

**Figure 4.**
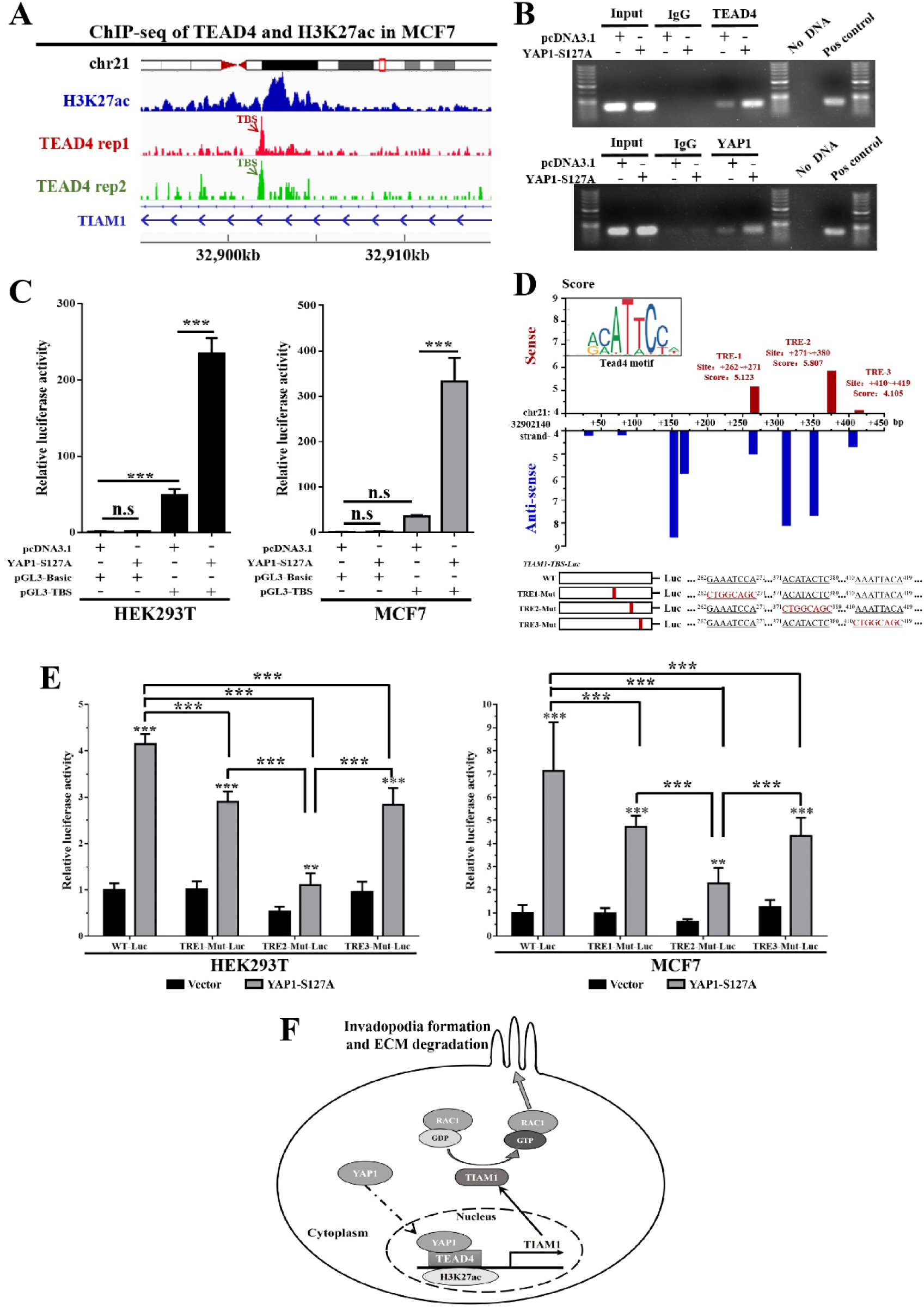
YAP1-TEAD4 transcriptionally activates TIAM1 expression through its enhancer region. (A) Diagram indicates that TEAD4 and H3K27ac peaks were located in similar regions of TIAM1 genome. Analysis was based on ChIP-seq data from ENCODE database (GSM1010860 and GSM945854). TEAD4-binding region was defined as the Tead4-binding site (TBS). (B) The YAP1-S127A mutant induced YAP1 and TEAD4 protein binding to the TBS region of the TIAM1 genome in MCF7 cells. Chromatin and proteins were cross-linked, and mouse monoclonal anti-TEAD4 and anti-YAP antibodies were used for the pull- down experiment. TBS of TIAM1 was amplified and measured by agarose gel electrophoresis or RT-qPCR (Supplemental Figure 6B). Mouse IgG was used as negative control, and histone H3 was used as positive control. (C) Dual luciferase reporter assay revealed that the TBS region exhibited transcription abilities and was significantly enhanced by the YAP1-S127A mutant both in HEK293T and MCF7 cell lines. ***p<0.01 by ANOVA. (D) JASPAR was used to identify potential TEAD4 regulation elements (TRE) in TBS. Three potential TREs were identified in the TBS region. (TRE-1: site 262∼271, sequence: GGAAATCCAT, score: 5.123; TRE-2: site 371∼380, sequence: AACATACTCC, score: 5.807, TRE-3: site 410∼419, sequence: AAAATTACAC, score: 4.105). According to the Tead4 motif form JASPAR, “CTGGCAGC” sequenced represented the low binding ability of TEAD4. Thus, this sequence was used in TRE mutation screening of TBS. (E) A dual luciferase reporter assay was used to identify the potential regulatory elements of the TBS activated by the YAP1-S127A mutant. TRE2 (+371∼+380) represented a significant transcription-enhancing ability in HEK293T and MCF7 cell lines. **p<0.01 and ***p<0.001. Student’s t test was used for intragroup comparisons, and ANOVA was used in intergroup comparisons. (F) Model for how the Hippo pathway regulates TIAM1 expression and induces invadopodia formation.

To identify the putative TEAD-response elements (TREs), we used the JASPAR program (jaspar.genereg.net) to search for the TEAD4 motif (as shown in Fig. S6D) in TBS region. Three potential TREs were identified in the positive-sense strand of the TBS region (TRE-1: 262-271, score: 5.123; TRE-2: 371-380, score: 5.807; TRE-3: 410-419, score: 4.105) (Fig. 4D and Fig. S6E). To determine the activity of each TREs, firstly, we analyzed the frequency matrix profile of TEAD4 from the JASPAR database (Fig. S6D). We found the sequence of “CTGGCAGC” showing low TEAD4 binding ability. Then, we performed a mutation scan to mutate each potential TREs sequences to “CTGGCAGC” (Fig. 4D). Dual luciferase assays showed that either TRE-1 or TRE-3 mutations could partially reversed YAP-S127A activated luciferase activity in MCF7 and HEK293T cells, whereas TRE-2 (371-+80) mutation could almost abrogated YAP-S127A regulated transcriptional activity (Fig. 4E). Therefore, TRE-2 is the most critical region within the enhancer of the TIAM1 genome. Taken together, our research elaborated a novel mechanism which YAP1-TEAD4 transcriptionally activate TIAM1 expression through its enhancer, and then increase RAC1 activity to induce invadopodia formation in breast cancer cells (Fig 4F).

## Discussion

In the current study, we demonstrated that YAP1, the key component of Hippo signaling, could induce invadopodia formation in breast cancer cells. Furthermore, we revealed that YAP1-TEAD4 could activate TIAM1 transcription in an enhancer dependent manner, which subsequently increases RAC1 activity and promotes invadopodia formation. These findings unveiled the novel role of YAP1 in invadopodia formation, and provide potential molecular targets for preventing tumor metastasis in breast cancer.

Invadopodia has been widely reported to induce tumor invasiveness and metastasis through degrading the extracellular matrix in various cancer (23). Invadopodia formation is a complex biological process and involves numerous signals, such as Src/Arg kinase, integrin-mediated signaling, EMT related pathway, Rac-GTPases mediated signaling, and matrix mechano-regulation (30–34). RAC1, a member of Rac-GTPase family, has been reported to play a critical role for promoting invadopodia formation (46, 48). Mechanically, activation of RAC1 has been demonstrated to be required for cortactin tyrosine phosphorylation which is a crucial step for invadopodia assembly (48). In the current study, we elucidated a novel function of YAP1 in invadopodia formation through increasing RAC1 activity. However, considering the promoting effect of the Hippo pathway in EMT signaling (19) and the presence of the SH3 domain in the YAP1 protein which exhibits potential catalytic activity for Src kinases (52, 53), we cannot completely exclude the other regulatory mechanisms between the Hippo pathway and invadopodia. Future studies are needed to further understanding of the mechanism regulating invadopodia through the Hippo signaling.

In current study, in order to clarify how YAP1 induced RAC1 activation, the bio-informatic analysis was performed, and a RAC1-specific guanine nucleotide exchange factor (GEF), TIAM1, was identified as a downstream effector of YAP1-TEAD complex. TIAM1 mediated the exchange of guanosine diphosphate (GDP) for guanosine triphosphate (GTP), and the binding of GTP induced a conformational change in RAC1, which activated RAC1 signaling pathways and promoted tumor metastasis (54). In addition, TIAM1 was also reported to affect other cellular processes, such as resistance to anoikis and induction of c-MYC induced apoptosis, via mechanisms that did not involve GEF activity (55, 56). Our findings demonstrated that YAP1 could up-regulate TIAM1 expression, and increased RAC1 activation and invadopodia formation. Interestingly, a recent study has reported that TIAM1 could promoted TAZ degradation in the cytoplasm and suppressed TAZ/YAP interaction with TEADs in the nucleus (57). Together with our findings, a negative feedback regulation could exist in the YAP1-TIAM1 axis and might play a role in maintaining balance between Hippo signaling and cytoskeleton dynamics.

The molecular mechanism of YAP1 inducing TIAM1 transcription is unknown. The conventional viewpoint is that YAP1 acts as a transcriptional coactivator via the promoter of downstream genes (58). However, in breast cancer, genome-wide analysis of YAP/TAZ-binding sites showed that nearly 91% YAP/TAZ-bound cis-regulatory regions coincide with enhancer elements, which located distant from transcription start sites (49). Enhancer appears to be an important regulatory element of YAP/TAZ-driven gene expression. In our study, we demonstrated that YAP1-TEAD4 activated TIAM1 transcription through binding to its enhancer region, rather than its promoter. Recently, the activation effects of YAP1 on enhancer were revealed by other studies. Zhu C, et al. demonstrated YAP1 functions as co-regulators of estrogen-regulated genes on enhancers in breast cancer (59). YAP/TAZ has also been reported to flag a large set of enhancers with super-enhancer-like functional properties (60). These studies unveiled an enhancer-mediate regulation of YAP1 protein, which might broaden our understanding of gene transcription in Hippo pathway.

In summary, our study demonstrated YAP1 could up-regulate TIAM1 expression through binding to its enhancer, which increased RAC1 activation and induced invadopodia formation in breast cancer. These findings reveal the role of Hippo signaling in invadopodia formation and provide potential molecular targets for preventing tumor metastasis in breast cancer.

## Materials and Methods

### Tissue arrays and Immunohistochemistry (IHC)

Human breast cancer tissue arrays (Shanghai Outdo Biotech Co. Ltd, Cat. #HBre-Duc060CD-01) were used to examine YAP1 expression in normal and cancer tissues. Slides were dewaxed, rehydrated and heated in sodium citrate buffer (0.01 M, pH 6.0) for antigen retrieval. Then, slides were rinsed with water and PBS with 0.1% Tween 20. Endogenous peroxidase was inhibited by incubation with freshly prepared 3% hydrogen peroxide with 0.1% sodium azide. Nonspecific staining was blocked through incubation in 5% bovine serum albumin for 2 hours. Next, slides were incubated overnight with YAP1 antibodies (Cell Signaling Technology, Cat. #4912) at a 1:100 dilution. Subsequently, after washing three times with PBS, the slide was incubated with a biotinylated secondary antibody for 2 hours. Immunostaining was performed using the DAB Horseradish Peroxidase Color Development Kit (Wuhan BosterBio Co. Ltd, Cat. #AR1022). Counterstaining was performed with hematoxylin. The results were analyzed under a microscope. In addition, scanning of this tissue array subject to hematoxylin-eosin staining was provided by the supplier (Shanghai Outdo Biotech Co. Ltd).

YAP1 expression and location were semiquantitatively measured via immunostaining scoring. YAP1 expression levels were calculated based on proportion score and intensity score. The proportion score reflected the fraction of positively stained cells (0, none; 1, ≤10%; 2, 10% to ≥25%; 3, >25% to 50%; 4, >50%), and the intensity score represented the staining intensity (0, no staining; 1, weak; 2, intermediate; 3, strong). Then, a total expression score was obtained by multiplying the proportion score and the intensity score, and YAP1 expression was categorized as level 1 (low grade, score 0–3), level 2 (medium grade, score 4-6) or level 3 (high grade, score greater than 6). The nuclear localization of YAP1 was calculated as the nucleus score. The nucleus score represented the fraction of positively stained nuclei (0, 0-10%; 1, 11-30%; 2, 31-70%; 3, 71-100%).

### Cell culture and transfections

MCF7 and HEK 293T cell lines were cultured in Dulbeccos modified Eagle’s medium (DMEM) (KeyGEN, Cat. #KGM12800N). MDA-MB-468 and MDA-MB-231 cell lines were cultured in Leibovitzs L -15 medium (L15) (KeyGEN, Cat. #KGM41300N). The T47D cell line was cultured in RPMI-1640 medium (1640) (KeyGEN, Cat. #KGM31800N). DMEM, L15 and 1640 culture media were supplemented with 10% fetal bovine serum (MULTICELL, Cat. #086-150) and 1% penicillin/streptomycin (KeyGEN, Cat. #KGY0023). The MCF10A cell line was cultured using an MEGM kit (Lonza/Clonetics Corporation, Cat. #CC-3150). All cells were cultured at 37°C in a 5% CO_2_ incubator. Plasmids pcDNA3.1-YAP1, pcDNA3.1-YAP1-S127A (Flag-tagged) and pcDNA3.1-YAP1-S94A (GFP-tagged) were gifted by Professor Bin Zhao, Zhejiang University. Small interfering RNAs (siRNAs) used for gene knockdown were provided by Guangzhou RiboBio Co. Ltd. The siRNA sequences are provided in Table S1. Cells were transfected with the indicated plasmids or siRNAs using Lipofectamine®2000 transfection reagent (Thermofisher, Cat. #11668019). Empty vector or non-targeting siRNA were used as negative controls. Biochemical and cell biological studies were performed 48 to 72 h after transfection.

### ECM degradation assay

ECM degradation assays were performed as previously described (61). Briefly, 12-mm coverslips were first incubated in 20% nitric acid for 2 hours and then washed in ddH_2_O for 4 hours. Then, coverslips were pretreated with 50 μg/mL poly-L-lysine (Sigma-Aldrich, Cat. #P8920) for 20 minutes. Coverslips were washed twice with PBS. Next, coverslips were cross-linked with 0.5% glutaraldehyde for 15 minutes and washed three times with PBS. Subsequently, each coverslip was inverted onto a 30- μl droplet of 1:8 Oregon Green™ 488 Conjugate Gelatin (Life Technologies, Cat. #G13186) with 0.1% porcine gelatin (Sigma, Cat. #G- 2500) for 10 min. Then, coverslips were incubated in 5 mg/mL sodium borohydride for 3 minutes followed by PBS washes. Coverslips were sterilized with 70% ethanol for 30 minutes after three PBS washes and incubated at 37℃ in complete growth medium for 1 hr. Twenty thousand cells were seeded on each coverslip, cultured for 24 hours, and processed for immunofluorescence. Alexa 594 phalloidin (Life Technologies, Cat. #A22287) was used to label ß-Actin at a 1:100 dilution, and DAPI was used for nuclear straining. Images were captured via a confocal laser-scanning microscope at 400x magnification. Gelatin degradation was calculated as the ratio of cells with degraded gelatin to the total cells per view field. Each experiment was performed in triplicate.

### Immunofluorescence

Twenty thousand cells were seeded on each coverslip, which was prepared using 0.1% porcine gelatin as described for the ECM degradation assay. After culture for 24 hours, cells were fixed in 4% paraformaldehyde at room temperature for 30 minutes and permeabilized with 0.1% Triton X- 100/PBS for 10 min. Nonspecific staining was blocked though incubation in 5% bovine serum albumin/PBS for 2 hours. Subsequently, cells were incubated at 4℃ with a cortactin antibody (Abcam, Cat. #ab33333) at a 1:100 dilution overnight and followed by 1:100 IFKine™ Green Donkey Anti-Mouse or Dylight 649 Goat Anti-Mouse secondary antibody (Abbkine, Cat. #A24211; A23610) and 1:100 phalloidin (Life Technologies, Cat. #A22287) for 2 hours. After washing, the nucleus was strained with DAPI, and coverslips were placed facedown onto a drop of anti-fading mounting medium on a microscope slide. Images were captured via confocal laser-scanning microscopy at 400x magnification. Each experiment was performed in triplicate.

### Transwell migration/invasion assay

Corning Transwell plates (24 wells, pore size 8 μm, Cat. #3422) were purchased for Transwell assays. For invasion experiments, Transwell filters were coated with 30 μl 1:8 diluted Matrigel (BD, Cat. #356234) on the upper surface of the polycarbonic membrane. Then, 1*10^5^ cells were harvested in 100 μl of serum-free culture medium and added to the upper compartment of the chamber. A total of 600 μl 30% fetal bovine serum medium was placed in the bottom compartment of the chamber as a chemoattractant. After 24 hours of culture at 37℃ with 5% CO_2_, the medium was removed, and the noninvaded cells on the upper side of the chamber were scraped off with a cotton swab. The migrated cells were fixed with 4% paraformaldehyde for 30 minutes and stained with 0.1% crystal violet/methanol for 15 minutes. After washing and drying, the number of migrated cells was counted in four randomly selected visual fields from the central and peripheral portion of the filter via an inverted microscope (100X magnification). Each assay was repeated three times.

### Wound-healing assay

Cells transfected with target siRNAs or plasmids were plated in each well of a 6-well culture plate at 90% cell density. On the next day, after cell adhesion, a scratch was created using a micropipette tip. The migration of cells towards the wound was monitored daily, and images were captured at 24-hr time intervals. Each assay was repeated three times.

### 3D tumor spheroid invasion assay

The method employed for 3D tumor spheroid invasion assays was previously described (40). Briefly, MCF7 cells were transfected with pcDNA3.1-YAP1 (G418 resistant) or empty vector as previously described and treated with 500 μg/ml G418 (Santa Cruz, Cat. #sc-29065) for 4 weeks to obtain stable cell lines. Then, cells were dilute to 1*10^4^ cells/ml and transferred to ultralow attachment 96-well plates (Corning, Cat. #7007) (200 μl per well). Cells were cultured at 37°C in a 5% CO_2_ incubator for 4 days to obtain tumor spheres. Next, 100 μl/well of growth medium was gently removed from the spheroid plates, and 100 μl/well Matrigel was added (BD, Cat. # 356234). The plates were transferred into a 37 °C incubator for 1 hour to allow the Matrigel to solidify, and 100 μl/well of complete growth culture medium was added. After 72 hours of culture, images were captured via an inverted microscope (200x magnification). Each assay was repeated three times.

### Microarray Gene Expression Profiling

Total RNA was collected from MCF7-Vector and MCF7-YAP1-S127A cells using TRIzol (Takara, Cat. #9108) reagent according to the manufacturer’s protocol. RNA was quantified using a NanoDrop ND-2000 (Thermo Scientific), and RNA integrity was assessed using an Agilent Bioanalyzer 2100 (Agilent Technologies). Sample labeling, microarray hybridization and washing were performed based on the manufacturer’s standard protocols. Briefly, total RNA was transcribed to double-stranded cDNA, synthesized into cRNA and labeled with cyanine-3-CTP. The labeled cRNAs were hybridized onto the microarray (Agilent SurePrint G3 Human Gene Expression v3 Panel). After washing, the arrays were scanned using an Agilent Scanner G2505C (Agilent Technologies).

Feature Extraction software (version10.7.1.1, Agilent Technologies) was used to analyze array images to obtain raw data. Genespring (version 13.1, Agilent Technologies) was employed to complete the basic analysis with raw data. Differentially expressed genes were then identified based on fold-change. The threshold set for up- and downregulated genes was a fold-change ≥ 1.5. Gene ontology analysis and Kyoto Encyclopedia of Genes and Genomes (KEGG) pathway enrichment of the differentially expressed genes were performed using DAVID software (https://david.ncifcrf.gov/tools.jsp).

### Quantitative real-time PCR

Total RNA was extracted with TRIzol (Takara, Cat. #9108) according to the manufacturer’s protocol. Reverse-transcription was performed using the PrimeScript^TM^RT Master Mix Reagent Kit (Takara, Cat. #RR036A) and quantitative real-time PCR was performed using the SYBR® Premix Ex Taq II Reagent Kit (Takara, Cat. Cat. #RR820A) according to the manufacturer’s protocol. Target genes were amplified using specific primers, and the human glyceraldehyde-3-phosphate dehydrogenase (GAPDH) gene was used as an endogenous control. Primer sequences used in this research are provided in Table S2. Data were analyzed using the comparative Ct method (2^-ΔΔCt^). Three separate bio-replications were performed for each experiment.

### Western blot assay

Total protein was extracted via RIPA lysis buffer with phenylmethylsulfonyl fluoride. Extracted proteins were boiled for 5 minutes in loading buffer before separation on 15% SDS-PAGE gels. After electrophoresis, separated protein bands were transferred into polyvinylidene fluoride membranes (Millipore, Cat. #IPVH00010). Then, membranes were blocked in 5% skim milk. Primary antibodies against YAP, TIAM1, TEAD4, FLAG, GFP, RAC1 and GAPDH were diluted according to the manufacturers’ instructions and incubated overnight at 4°C. Manufacturer names and catalog numbers of the antibodies used in this research are provided in the Table S3. Horseradish peroxidase-linked secondary antibodies were added at a dilution ratio of 1:1000 and incubated at room temperature for 2 hours. The membranes were washed with TBST buffer three times and visualized using the ECL Kit (Thermofisher, Cat. #34096).

### GTP-bound GTPase pull-down assay

GTP-bound GTPase pull-down assays were performed using the Active Rac1/CDC42 Detection Kit (Cell Signaling Technology, Cat. #8815, 8819). After transfection or verteporfin treatment, MCF7 cells were harvested according to the manufacturers instruction. Enrichment of active GTP - bound GTPase was performed using GST-PAK1-PBD fusion protein beads. Then, the proteins on the beads or total cell lysates were extracted and tested via Western blot. The full details of the pull-down procedure are described in the manufacturers instruction.

### Chromatin Immunoprecipitation (ChIP)

ChIP was performed using the SimpleChIP Enzymatic Chromatin IP Kit (Cell Signaling Technology, Cat. #9003). MCF7 cells were transfected with empty vector or YAP1-S127A for 48 hours. After chromatin and proteins were cross-linked, 500 μl diluted cross-linked chromatin was incubated overnight with 5 μg mouse monoclonal anti-TEAD4 antibody (Abcam, Cat. #ab58310), 5 μg mouse monoclonal anti-YAP antibody (Cell Signaling Technology, Cat. #14074) or 1 μg normal mouse IgG (Cell Signaling Technology, Cat. #2729). Histone H3 (D2B12) XP® Rabbit mAb (Cell Signaling Technology, Cat. #4620) was used as a positive control. The TEAD4 binding site and positive control gene PRL30 were quantified by PCR and real-time quantitative PCR methods. The TEAD4 binding site was determined based on the ENCODE dataset GSM1010860 (HG19, Chr 21: 32901686-32902140, strand: minus), and the primer sequences were as follows: Forward: AAGAAGATACATTGAAGAGATAA;

Reverse: TGAGTTCCAAGCATAAGA.

PRL30 primers were provided in the kit. The full details of the ChIP procedure are described in the ChIP kit’s instructions.

### Plasmid construction and Site-directed mutagenesis

The TEAD4-binding site (TBS) of the TIAM1 genome in MCF7 cells was determined by the ENCODE project (HG19, Chr 21: 32901686-32902140, strand: minus) and synthesized by TsingKe BioTech Co. Ltd. The TBS sequence was cloned into the pGL3-Basic vector (Promega, Cat. #E1751) using BglII/HindIII and named pGL3-TBS. Site-directed mutagenesis of pGL3-TBS was performed using the Trelief™ SoSoo Cloning Kit (TsingKe, Cat. #TSV-S1) according to the manufacturer’s protocol. The following primers were used for mutagenesis:

TRE1-Mut:

#### Forward

5‛-ATCAGCAAAGCTGCCAGCTTTCAGGGAGTTTAAGACCTTG-3‛,

#### Reverse

5‛-AAGCTGGCAGCTTTGCTGATAAAGAAGATACATTGAAGAG-3‛

TRE2-Mut:

#### Forward

5‛-GCTGCCAGTCCCTTCAGTAGCTCTTGAATACCTTGAGT-3‛,

#### Reverse

5‛-CTACTGAAGGGACTGGCAGCCTCGTTTCTGAAATCTTATGC-3‛. TRE3-Mut:

#### Forward

5‛-AGGCTGCCAGTGAGTTCCAAGCATAAGATTTCAGAAACGAGG-3‛

Reverse:

5‛-GCTTGGAACTCACTGGCAGCCTAGTAATTGTTACAGT-3‛.

### Dual luciferase assays

Briefly, 100 ng pGL3-Basic luciferase reporter plasmid (Promega, Cat. #E1751) with target sequence inserts were cotransfected into HEK293T and MCF7 cell lines with 200 ng of Control/YAP1-S127A construct and 10 ng of Renilla luciferase pRL-TK plasmid (Promega, Cat. #E2241) using Lipofectamine®2000 transfection reagent (Thermofisher, Cat. # 11668019). After 48 hours, dual luciferase assays were performed using the Dual-Luciferase® Reporter Assay System (Promega, Cat. #E1910). Luciferase activity was measured as the ratio of firefly luciferase signal to Renilla luciferase signal. All measurements were normalized to the control group.

### Public database and bioinformatics analysis

The gene expression profiles of purified tumor cells from 14 primary breast tumor tissues and 6 metastatic lymph nodes were downloaded from the Gene Expression Omnibus (GEO) database (GSE30480, (62)) and analyzed via Gene Set Enrichment Analysis (GSEA) software (http://software.broadinstitute.org/gsea/) (63). Clinical data and expression profiles of 252 breast cancer patients (GSE21653, (64)) and 508 breast cancer patients (GSE25066, (65)) were employed for Kaplan-Meier analysis using the R2: Genomics Analysis and Visualization Platform (http://r2.amc.nl). Prognosis analysis of TCGA breast invasive carcinoma dataset was performed using the SurvExpress program (http://bioinformatica.mty.itesm.mx:8080/Biomatec/SurvivaX.jsp) (37). The expression level of the TEAD family in breast cancer was determined by the cBioPortal program (www.cbioportal.org) using the Breast Cancer (METABRIC, Nature 2012 & Nat Commun 2016) dataset (samples=2509) (66, 67). Gene correlation analysis was performed using the R2: Genomics Analysis and Visualization Platform (http://r2.amc.nl) with the Auckland breast cancer dataset (GSE36771, (68)) and TCGA breast cancer dataset. TEAD4 and H3K27ac ChIP-seq data from MCF7 cells were downloaded from the ENCODE project (http://genome.ucsc.edu/ENCODE/downloads.html) (GSM1010860, GSM945854) and visualized via Integrative Genomics Viewer (IGV) software (http://software.broadinstitute.org/software/igv/) (69). Protein interaction analysis was based on the String database (http://www.string-db.org/) (70). TEAD4 motif analysis was performed using JASPAR software (http://jaspar.genereg.net/) (71).

### Statistical analysis

Statistical analysis was performed using the software package SPSS (version 19.0 for Windows; IBM, USA) and GraphPad Prism (version 6.01 for Windows; GraphPad Software, Inc., USA). All continuous data were presented as the mean ± SD. Continuous data were statistically analyzed via Student’s t-test (two-tailed) and analysis of variance (ANOVA), whereas categorical variables were compared using the χ2 test and Fisher test. Statistical significance was set at p < 0.05.

## Supporting information

Supplemental informatin

## Notes

### Competing Interest Statement

The authors have declared no competing interest.

