## Supplemental informatin for "YAP1 induces invadopodia formation by transcriptionally activating TIAM1 through its enhancer in breast cancer"

### YAP1-TEAD4 induces TIAM1 expression to promote invadopodia formation in breast cancer

**This PDF file includes:**

- Figures S1 to S6
- Tables S1 to S3
- Legends for Datasets S1 to S4

**Other supplementary materials for this manuscript include the following:**

- Datasets S1 to S4

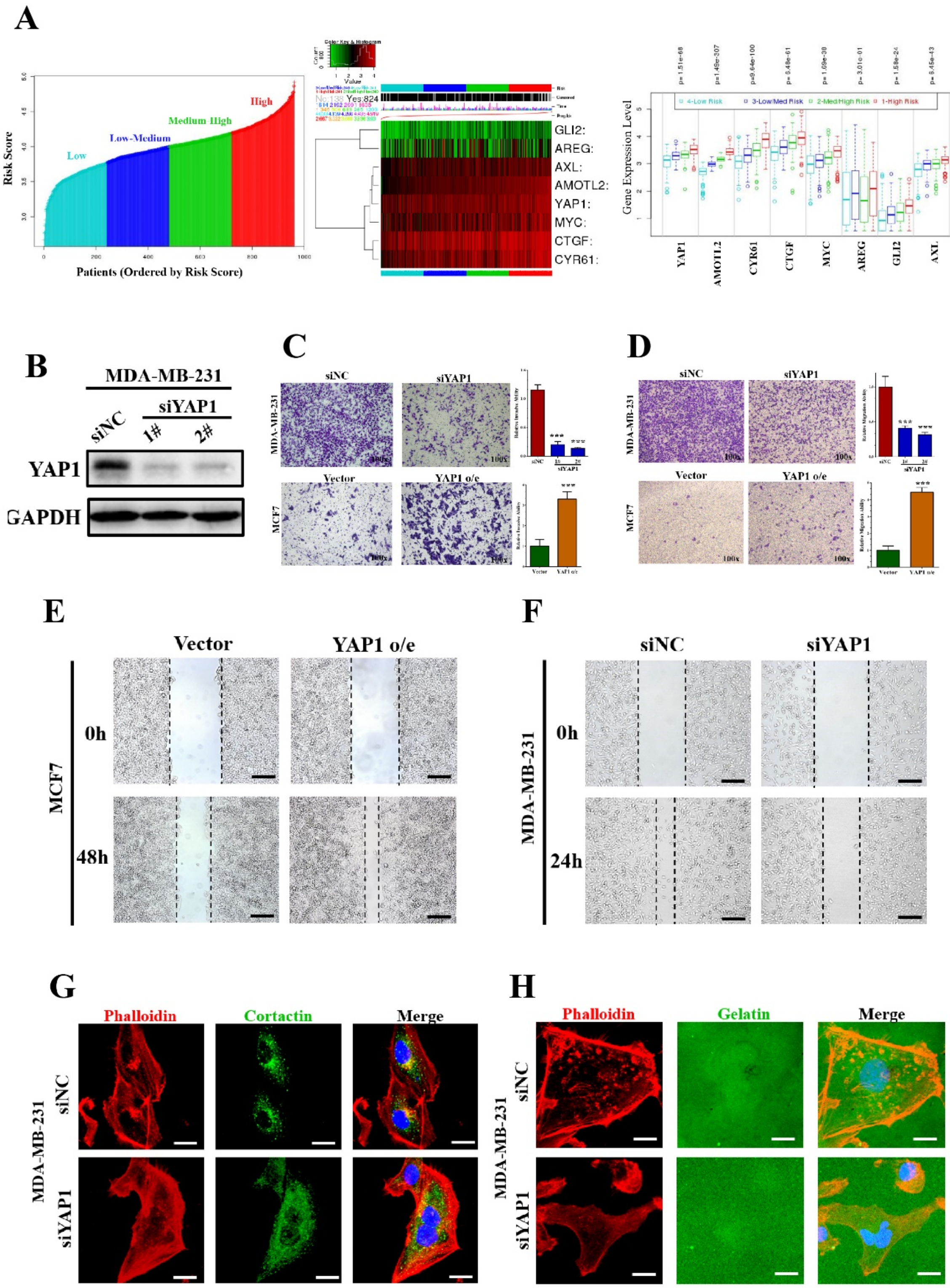


**Fig. S1. YAP1 is associated with tumor invasiveness and induces invadopodia formation in breast cancer.**

(A) According to the prognostic condition, IBC patients from TCGA dataset were ranked evenly into four groups (Low, Low-Medium, Medium-High, and High) (left) by risk score. Heat map (middle) and histogram (right) summarizing YAP1 and its target genes (AMOTL2, CYR61, CTGF, MYC, GLI2, AREG and AXL) expression in these IBC patients (ordered by risk score). This prognostic analysis was performed by SurvExpress program. (B) Western blot verified knockdown of endogenous YAP1 in MDA-MB-231 cells. (C) YAP1 expression promoted MDA-MB-231 and MCF7 cell migration. Transwell migration assay was performed, and representative images are presented (***p<0.001 by Student`s t test). (D) YAP1 expression promoted MDA-MB-231 and MCF7 cell invasion. Matrigel invasion assay was performed, and representative images are presented (***p<0.001 by Student’s t test). (E) YAP1 overexpression promoted cell migration in MCF7 cells. Wound-healing assay was performed, and representative images are presented (Scale bar: 200 μm). (F) YAP1 knockdown in MDA-MB-231 cells inhibited migration. Wound-healing assay was performed, and representative images are presented (Scale bar: 200 μm). (G) YAP1 knockdown in MDA-MB-231 cells reduced invadopodia formation. Scale bar: 20 μm. (H) YAP1 knockdown in MDA-MB-231 cells decreased ECM degradation ability (Scale bar: 20 μm).

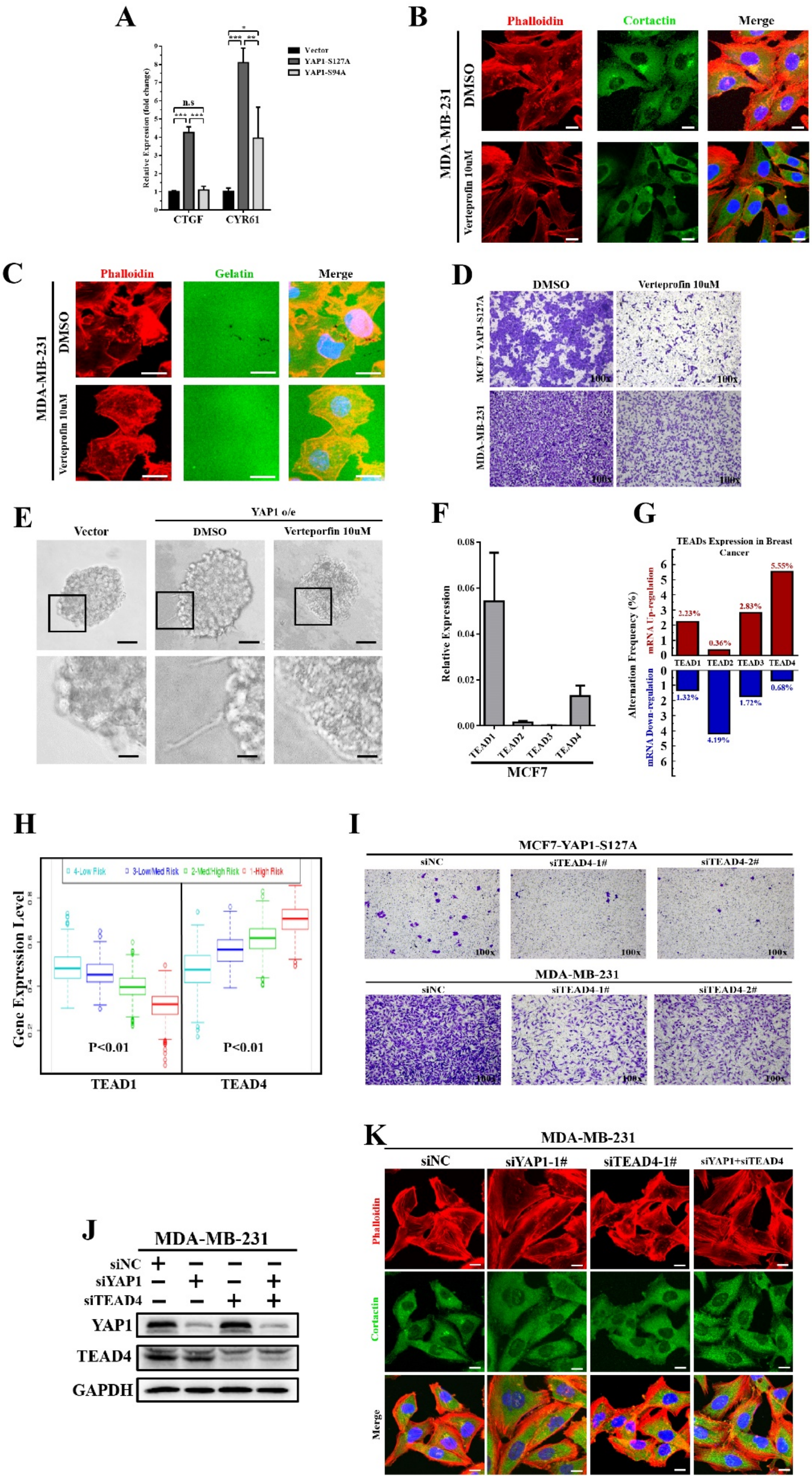

**Fig. S2. YAP1-TEAD4 interaction is essential for invadopodia formation and tumor migration.**

(A) mRNA expression levels of YAP1 target gene (CTGF and CYR61) in MCF7-Vector, MCF7-YAP1-S127A and MCF7-YAP1-S94A cells. GAPDH was used as an internal control. *$p<0.05$, **$p<0.01$ and ***$p<0.001$ by ANOVA. (B) Verteporfin inhibited invadopodia formation in MDA-MB-231 cells. Cells were seeded on 0.1% gelatin and treated with 10 μM verteporfin (DMSO was used as negative control). After 48 h of culture, invadopodia were visualized by colocalization of cortactin (green) and F-actin (red). Scale bar: 20 μm. (C) Verteporfin inhibited ECM degradation in MDA-MB-231 cells. MDA-MB-231 cells were seeded on Alexa Fluor 488-conjugated gelatin (green) and treated with 10 μM verteporfin (DMSO was used as negative control) for 24 hr. F-actin was stained with phalloidin (red), and nuclei were stained with DAPI (blue). The area of gelatin degradation appears as black area beneath the cells (scale bar: 20 um). (D) Transwell migration assay revealed that verteporfin decreased MCF7-YAP1-S127A and MDA-MB-231 cell migration ability. (E) 3D tumor sphere invasion assays revealed that verteporfin reduced pseudopod-like structure formation in MCF7-YAP1 cells (Scale bar: 40 um). (F) TEAD1-4 expression levels in MCF7 cells measured by RT-qPCR. GAPDH was used as an internal control. (G) Overview of expression of the TEAD family (TEAD1, TEAD2, TEAD3 and TEAD4) in breast cancer. TEAD4 exhibited the highest upregulation among the TEAD proteins. Analysis was based on the cBioPortal program using the Breast Cancer (METABRIC, Nature 2012 & Nat Commun 2016) dataset (n=2509). (H) SurvExpress analysis revealed that TEAD4 exhibited a positive correlation with poor prognosis in breast cancer ("Breast cancer recurrence data, 9 datasets from 7 authors" database was used in the analysis). (I) Knockdown of TEAD4 inhibited cell migration in MCF7-YAP1-S127A and MDA-MB-231. Transwell migration assay was performed, and representative images are presented. (J) Knockdown of endogenous YAP1 and/or TEAD4 expression in MDA-MB-231 cells using siRNAs. (K) Knockdown of endogenous YAP1 and/or TEAD4 expression reduced invadopodia formation in MDA-MB-231 cells. Cortactin (green) and F-actin (red). Scale bar: 20 μm.

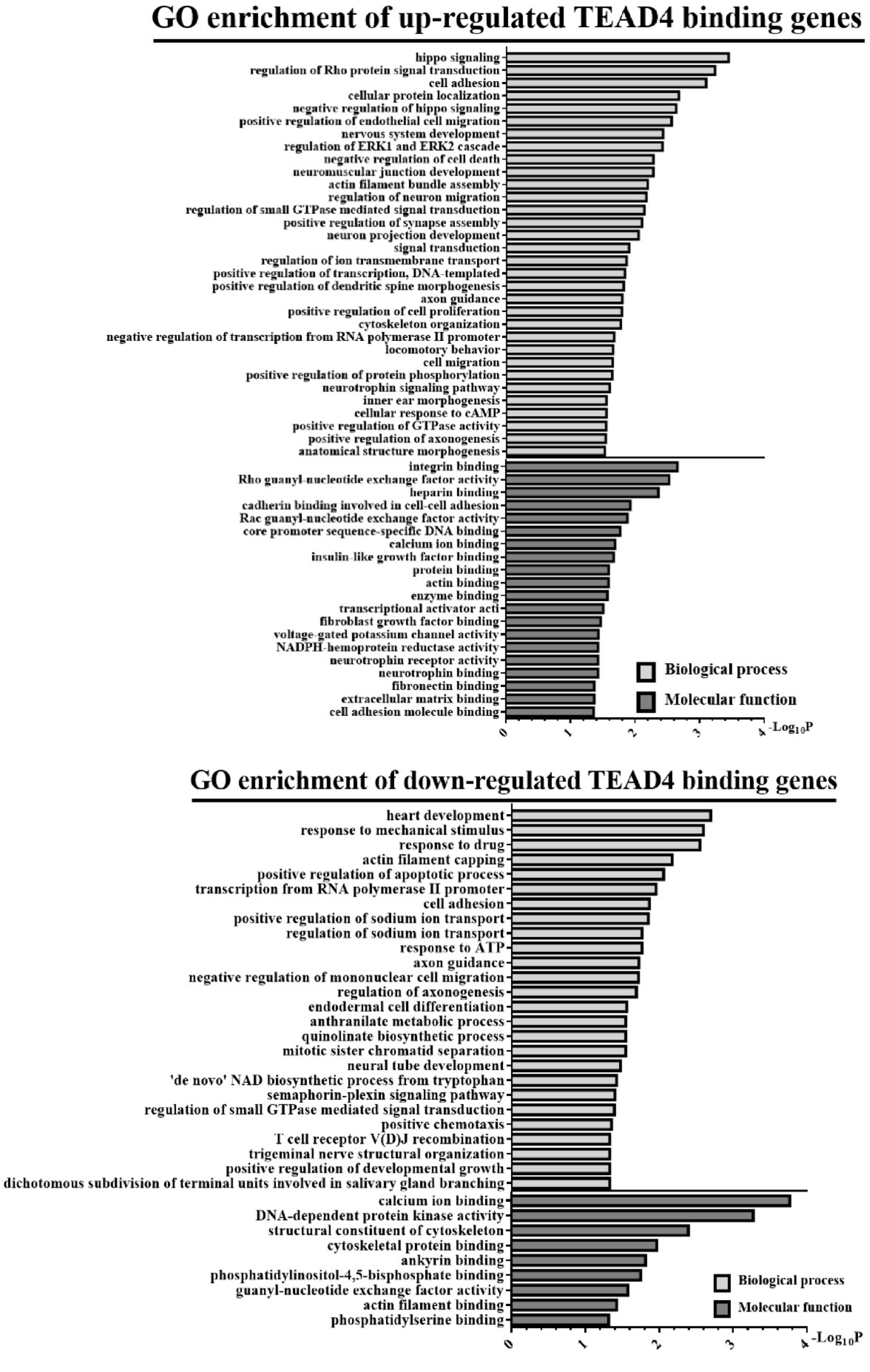


**Fig. S3. Gene Ontology enrichment analysis of the up/downregulated TEAD4 binding genes affected by the YAP1-S127A mutant in MCF7 cell line.**

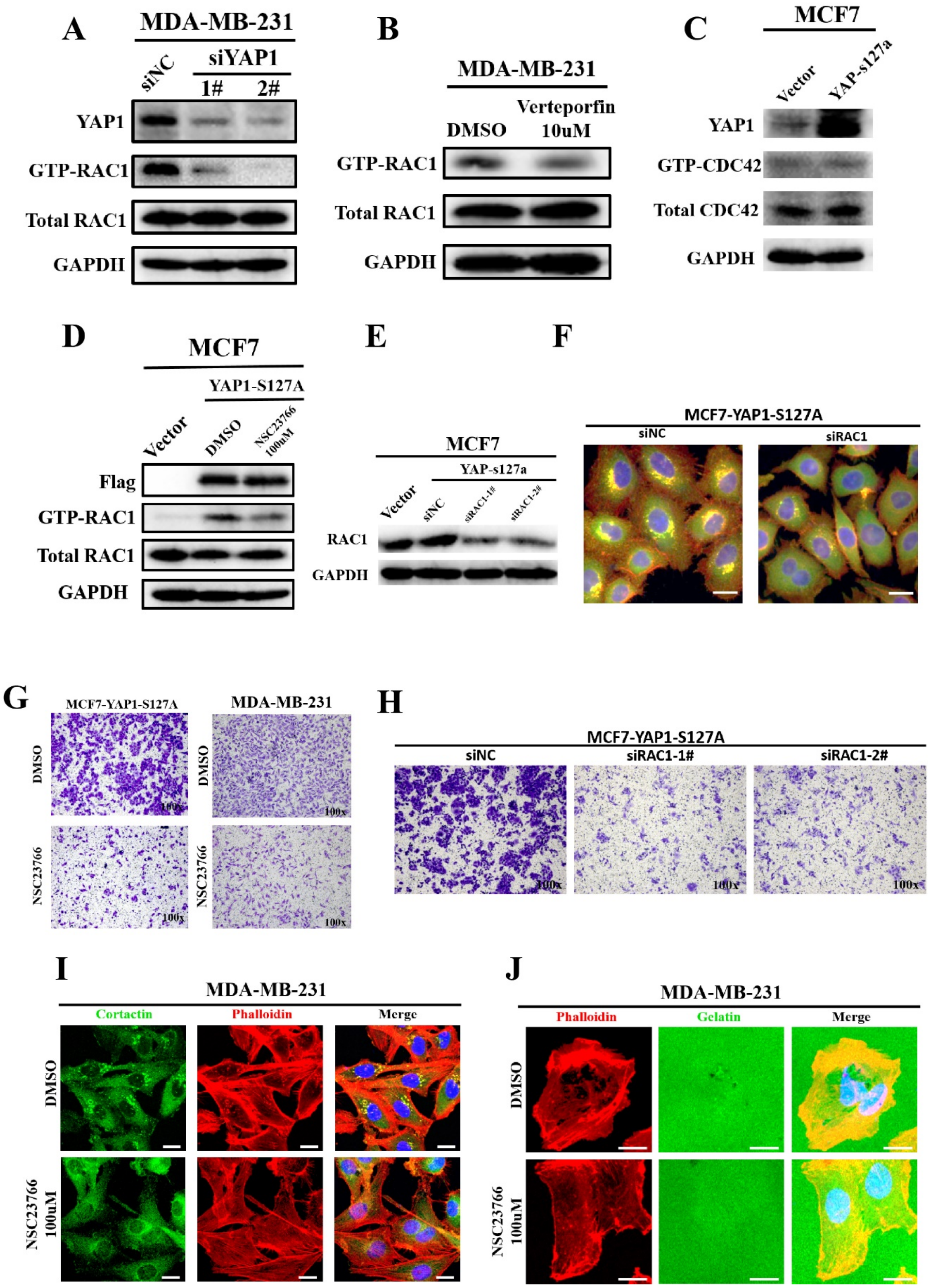


**Fig. S4. YAP1 induces invadopodia formation in a RAC1-dependent manner.**

(A) As shown in GTP-bound GTPase pull-down experiments and Western blots, knockdown of YAP1 expression reduced RAC1 activity in MDA-MB-231 cells. (B) Verteporfin reduced RAC1 activity in MDA-MB-231 cells. MDA-MB-231 was treated with 10 μM verteporfin (DMSO was used as a negative control) for 24 hr. Then, Rac1 activation was measured via GTP-bound

GTPase pull-down experiments and Western blots. (C) GTP-bound GTPase pull-down experiment demonstrated that YAP1-S127A overexpression did not affect CDC42 activity in MCF7 cells. (D) Using NSC23766 to inhibit RAC1 activation in MCF7-YAP1-S127A cells. MCF7 cells (transfected with the flag-tagged-YAP1-S127A overexpression plasmid) were treated with 100 μM NSC23766 (DMSO was used as a negative control) for 12 hr. Then, Rac1 activation was measured by GTP-bound GTPase pull-down experiments and Western blots. (E) Knockdown of RAC1 expression in MCF7 cells transfected with the YAP1-S127A plasmid. (F) Knockdown of RAC1 expression inhibited invadopodia formation in MCF7-YAP1-S127A cells (Green: cortactin; Red: F-actin; Blue: DAPI, Scale bar: 20 um). (G) NSC23766 inhibited MCF7-YAP1-S127A and MDA-MB-231 cell migration at a dose of 100 μM. Transwell migration assay was performed, and representative images are presented. (H) Knockdown of RAC1 expression reduced MCF7-YAP1-S127A cell migration. (I) After treatment with 100 μM NSC23766 for 24h, invadopodia formation was significantly reduced in MDA-MB-231 cells (Scale bar: 20 um). (J) After treatment with 100 μM NSC23766 for 24 h, ECM degradation was significantly reduced in MDA-MB-231 cells (Scale bar: 20 um).

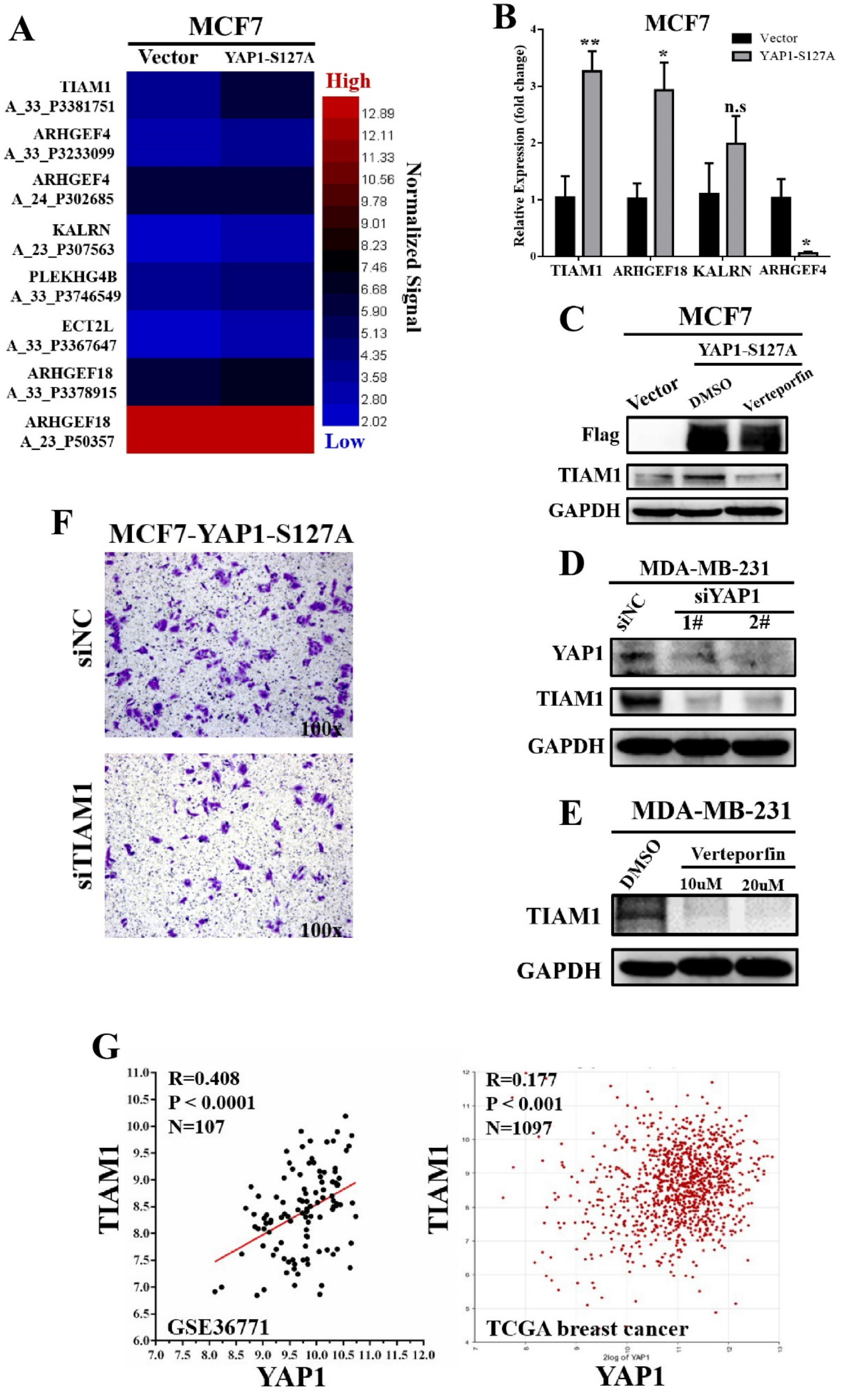


**Fig. S5. YAP1-TEAD4 up-regulates TIAM1 expression.**

(A) The heat map summarizes the upregulated TEAD4-binding genes that are attributed to

Rho/RAC guanyl-nucleotide exchange factor activity categories (GO: 0005089 and GO: 0030676) in MCF7 cells transfected with YAP1-S127A mutant (Color scale denoted normalized mRNA expression signal). (B) RT-qPCR verified mRNA expression fold-changes of the upregulated RAC1 associated TEAD4-binding genes in MCF7 cells transfected with YAP1-S127A mutant. *$p<0.05$ and **$p<0.01$ by Student's t test. (C) Verteporfin reversed YAP1-S127A induced TIAM1 expression at a dose of 10 μM in MCF7 cells as determined by Western blot. (D) YAP1 knockdown in MDA-MB-231 cells downregulated TIAM1 expression as determined by Western blot. (E) Verteporfin inhibited TIAM1 expression in MDA-MB-231 cells as determined by Western blot. (F) Knockdown of TIAM1 expression reduced migration in MCF7 cells transfected with the YAP1-S127A mutant. Transwell migration assay was performed, and representative images are presented. (G) TIAM1 expression was positively associated with YAP1 expression in clinical breast cancer specimens. Gene correlation analysis was based on 107 breast cancer patients in Auckland (GSE36771) (Left) and 1097 breast cancer patients from TCGA (Right). TCGA dataset analysis was based on R2: Genomics Analysis and Visualization Platform.

**A**

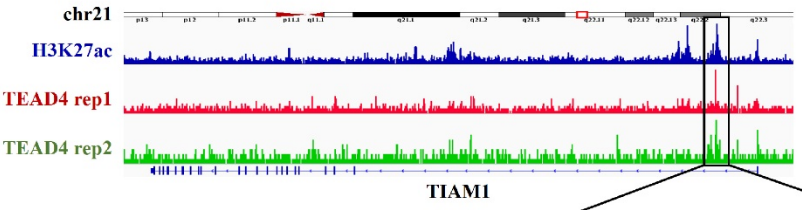


| | Range |
|---|---|
| TEAD4-rep1 peak | chr21:32901707-32902140 |
| TEAD4-rep2 peak | chr21:32901686-32902079 |
| **TBS region** | **chr21:32901686-32902140** |

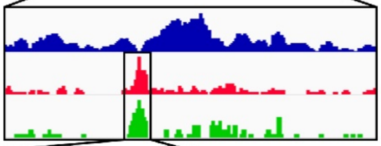

**TBS region:** hg19_dna range=chr21:32901686-32902140; strand: minus; 5`->3`
CCTTCCCTCGAGAGGAAGGGGGCATGAGGTGAATTTTCTCTGGAGTTAGGAGAGATGATACCAGGTTGGAGTTC
ATGGACTTTAAGTTTATAAACACATGTCTGCACAGCTGTCTTGCAATGACAGTTGCTCTCTGCATTTGAGGAGCA
GGTATTTGGGAAGGAGCAATGCTGGATGGGTGACCTAATTTATTTCCTTTGTTTCCTTCTGGCCTGAACTGGGCA
GGTAGCCCTGACCCACAAGGTCTTAAACTCCCTGAAAGGAAATCCATTTGCTGATAAAGAAGATACATTGAAGA
GATAAAGAAGAGAATGTTTGGTTTCCTGCTTCCTTCTCTGTTTACTCAAGGTATTCAAGAGCTACTGAAGGGAAC
ATACTCCTCGTTTCTGAAATCTTATGCTTGGAACTCAAAATTACACTAGTAATTGTTACAGTAGTAACAACCCATA
A

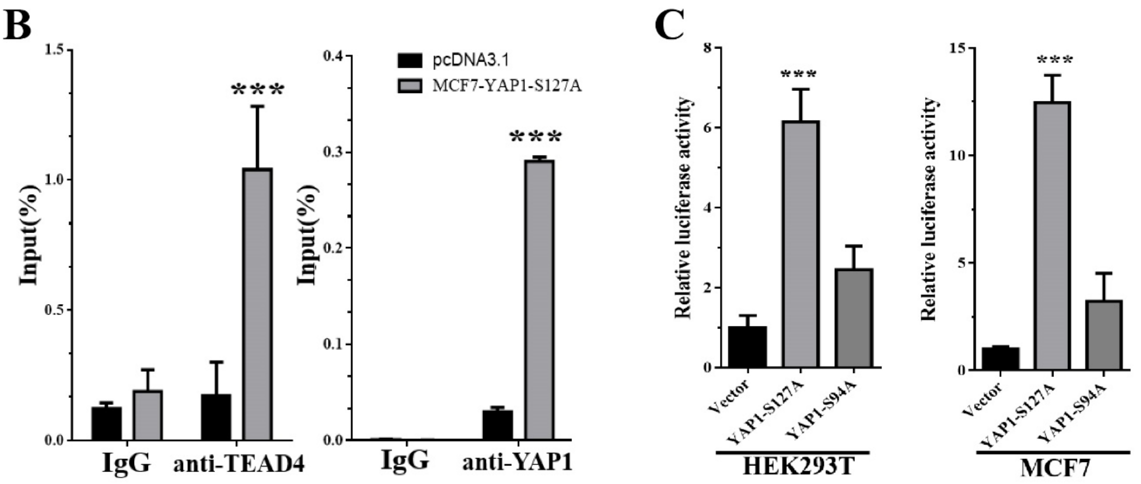


**D**

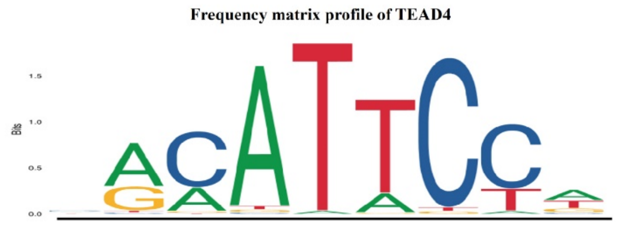


| Pos | 2 | 3 | 4 | 5 | 6 | 7 | 8 | 9 |
|---|---|---|---|---|---|---|---|---|
| A | 645 | 434 | 1011 | 21 | 213 | 0 | 49 | 572 |
| C | 22 | 1011 | 12 | 0 | 0 | 1011 | 1011 | 82 |
| G | 366 | 34 | 21 | 0 | 16 | 19 | 1 | 226 |
| T | 30 | 18 | 26 | 1011 | 1011 | 27 | 348 | 439 |

**E**

| Site | Strand | Sequence(5`-3`) | Score |
|---|---|---|---|
| 28~37 | Anti-sense | AAAATTCACC | 4.174 |
| 75~84 | Anti-sense | TAAAGTCCAT | 4.138 |
| 148~157 | Anti-sense | CAAATACCTG | 8.588 |
| 163~172 | Anti-sense | AGCATTGCTC | 5.818 |
| 260~269 | Anti-sense | GGATTTCCTT | 4.973 |
| 262~271 | Sense | GGAAATCCAT | 5.123 |
| 307~316 | Anti-sense | AACATTCTCT | 8.081 |
| 346~355 | Anti-sense | TGAATACCTT | 7.654 |
| 371~380 | Sense | AACATACTCC | 5.807 |
| 401~410 | Anti-sense | TGAGTTCCAA | 4.665 |
| 410~419 | Sense | AAAATTACAC | 4.105 |

**Fig. S6. YAP1-TEAD4 transcriptionally activates TIAM1 expression through its enhancer region.**

(A) IGV analysis of ChIP-seq data from the ENCODE database (GSM1010860 and

GSM945854). The TBS region is calculated as the aggregate of the TEAD4 binding peaks from two bio-replicates of ChIP-seq experiments. The TBS sequence is presented in the figure. (B) YAP1/TEAD4 binding region of TIAM1 genome in MCF7 cell line is obtained via ChIP and quantified using qPCR. ***p<0.001 by Student's t test. (C) Dual luciferase reporter assay revealed that compared with the YAP1-S94A mutant, YAP1-S127A could significantly increase luciferase activities in HEK293T and MCF7 cell lines co-transfected with pGL3-TBS and Renilla plasmids. ***p<0.001 by ANOVA. (D) Frequency matrix profile of TEAD4. (E) Detailed information of potential Tead4 regulation elements (TRE) in TBS.

**Table S1.** Target sequences of siRNAs.

**Target sequences of siRNAs**

| Name | Target Sequence (5`-3`) |
|---|---|
| si-TEAD4-1# | GCTTGTGGATGAAGTTGAT |
| si-TEAD4-2# | GGACACTACTCTTACCGCA |
| si-TIAM1-1# | GGAGCTGATTTGCAAGACA |
| si-TIAM1-2# | GCAGTTTGCATGAGATGAA |
| si-YAP1-1# | GCGTAGCCAGTTACCAACA |
| si-YAP1-2# | GGTGATACTATCAACCAAA |
| si-RAC1-1# | TGAAGAAGAGGAAGAGAAA |
| si-RAC1-2# | GGAACTAAACTTGATCTTA |

**Table S2.** Primer sequences.

| Primers` sequences | |
|---|---|
| **Primer** | **Sequence (5`-3`)** |
| GAPDH-F | CTCCTGCACCACCAACTGCT |
| GAPDH-R | GGGCCATCCACAGTCTTCTG |
| CTGF-F | AGGAGTGGGTGTGTGACGA |
| CTGF-R | CCAGGCAGTTGGCTCTAATC |
| CYR61-F | CCTTGTGGACAGCCAGTGTA |
| CYR61-R | ACTTGGGCCGGTATTTCTTC |
| YAP1-F | TAGCCCTGCGTAGCCAGTTA |
| YAP1-R | TCATGCTTAGTCCACTGTCTGT |
| TEAD4-F | AGTCAGGCACTGGACAAGC |
| TEAD4-R | GCTGGAGACCTGCTTCCTG |
| TIAM1_F | GATCCACAGGAACTCCGAAGT |
| TIAM1_R | GCTCCCGAAGTCTTCTAGGGT |
| ARHGEF4_F | ACCATCTTCGGGAACATCGAG |
| ARHGEF4_R | CTGGAAGTCGGCTTGATGCT |
| KALRN_F | TATCTCTGGTATCTCCGCTTGC |
| KALRN_R | CCTCGCTTATCACGACCCC |
| ARHGEF18_F | ATGACGGTCTCTCAGAAAGGG |
| ARHGEF18_R | GTGATTGGTCCGAGTTGCGT |

**Table S3.** List of antibodies.

#### Cat No. of the antibodies

| Antibody | Manufacture | Cat. No. |
|---|---|---|
| YAP | Cell signaling technology | 4912 |
| YAP (ChIP grade) | Cell signaling technology | 4074 |
| TIAM1 | Abcam | ab211518 |
| TEAD4 | Abcam | ab58310 |
| FLAG | Santa Cruz | sc-166355 |
| GFP | Santa Cruz | sc-81045 |
| GAPDH | Santa Cruz | sc-32233 |
| RAC1 | Cell signaling technology | 8631 |
| Cortactin | Abcam | ab33333 |
| mouse IgG | Cell signaling technology | 2729 |
| Histone H3 | Cell Signaling Technology | 4620 |

**Dataset S1 (separate file).** General characteristics of clinical specimens in tissue chip.

**Dataset S2 (separate file).** Gene set enrichment analysis of C6_Oncogenic signatures in GSE30480 dataset.

**Dataset S3 (separate file).** Different expression genes in MCF7 cells overexpressing YAP1-S127A vs. control plasmid.

**Dataset S4 (separate file).** Annotation of TEAD4 ChIP sequence data in MCF7 cell line from ENCODE database (GSM1010860) via ChIP-Seek software.
